# Antibacterial Action of Gliotoxin Realized Through Interaction with Tiol-containing Proteins

**DOI:** 10.1101/2022.08.23.504972

**Authors:** Alexey S. Vasilchenko, Elena V. Gurina, Konstantin A. Drozdov, Dmitry Andreev, Egor Lebedev, Anastasia V. Vasilchenko

## Abstract

Gliotoxin is a secondary metabolite of various fungi belonging to the class of epipolythiodioxopiperazines. The mechanism of gliotoxin cytotoxic activity on eukaryotes is established, while the precise interaction between gliotoxin and bacteria has not been clarified yet. The aim of this study was evaluating the gliotoxin action on model Gram-positive and Gram-negative bacteria. It was found that gliotoxin uptake rate was higher in *S. epidermidis* than *E.coli*. However, for Gram-negative bacteria, the bactericidal effect is achieved at higher doses of gliotoxin (10-100 μg/ml), compared to Gram-positive ones (0.75-6.25 μg/ml). Bactericidal effect had developed in 4 hours, which was the same for both *E. coli* MG1655 and *S. epidermidis*. Novel chip-based bioluminescent sensor with gel-immobilized *E.coli* MG1655 pKatG-lux revealed oxidative stress. However, pre-incubation of bacteria with Trolox did not affect the bactericidal properties of gliotoxin, while 2-merkaptoethanol and reduced glutathione significantly reduced gliotoxin’s bactericidal property. Another bioluminescent sensor *E.coli* MG1655 pIbpA-lux revealed heat shock stress in bacteria treated with gliotoxin. At the cellular level, exposure bacteria with gliotoxin accompanied by disruption of bacterial membranes.

## Introduction

Fungi are capable to produce secondary metabolites which is not vital for producers, but allow them to implement their life strategies. Among these metabolites antibiotics and mycotoxins have important place. Gliotoxin is one of the most controversial secondary metabolites produced by some fungi of the *Gliocladium* and *Aspergillus* species. Over the past 85 years since the discovery of gliotoxin [1], its status is still unclear. On the one hand, the antimicrobial properties of gliotoxin-producing strains of *Trichoderma virens* have found their application as the plant-protection agents (for example, SoilGard™) [2], as well as gliotoxin have potential as a drug with an oncotarget effect [3]. On the other hand, gliotoxin is considered as the pathogenicity factor of aspergillus and is involved in the development of immunosuppressive states of the human [4,5].

Thus, understanding the role of gliotoxin in fungal ecology and its biotechnological importance requires more data on its action against prokaryotes.

Gliotoxin (GTX) by its physicochemical properties belongs to the class epipolythiodioxopiperazine (ETPs), which contain an intra- or intermolecular sulfur bridge and a diketopiperazine (DKP) core [6]. To date, it is known at least 20 different epidiathiodioxopiperazines produced exclusively by fungi of marine and soil habitats [7].

According to modern concepts, the presence of a disulfide bridge in the gliotoxin molecule determines the specificity of its action on eukaryotic cells. One of the postulated modes of GTX action is producing of reactive oxygen species. In the presence of a reducing agent, the disulfide bond in the GTX molecule is reduced to dithiols, and then auto-oxidize back to disulfides, what accompanies converting oxygen into superoxide radical [8, 9]. Generated ROS causing single- and double-stranded DNA breaks [8].

Alternatively, or simultaneously, some ETPs can form disulfide bridges with protein cysteine thiols, what led to inactivation of various enzymes which containing cysteines in their active site [10]. Thus, generation of reactive oxygen species which affect DNA, lipids and proteins, and inactivation of cellular thiol-containing enzymes suggested as the most likely mechanisms of ETPs toxicity.

However, no clear which kind of stated mechanisms work in the gliotoxin-prokaryote interaction. There is still no clear understanding of either bactericidal concentrations in relation to bacteria with various types of cell wall architecture.

In this work, we studied bactericidal effect of gliotoxin on model Gram-positive and Gram-negative bacteria. Using bacterial biosensor strains, we had shown *in vivo* whether the toxic effect of gliotoxin depends on the generation of reactive oxygen species or disturbance of protein structure.

## Materials and Methods

### Gliotoxin biosynthesis and purification

Gliotoxin was obtained by cultivation of *Aspergillus fumigatus* UTMN1 on liquid Weindling broth (sucrose – 15 g/L, (NH_4_)SO_4_ – 1.6 g/L, K_2_HPO_4_ – 0.83 g/L, MgSO_4_ – 0.41 g/L, FeCl_3_ – 0.008 g/L, peptone – 0.01 g/L, pH=3.3-3.8) at 28 °C, rpm 100 for 4 days. Then fungal micellium was pellet by centrifugation and filtered through PVDF-membrane (WVR, US). Obtained culture medium mixed with chloroform (2:1 v/v). Separation of chloroform from the culture liquid carried out with separating funnel. Extraction was performed three times.

Next, chloroform was evaporated on a rotary evaporator to obtain a dry residue, which was re-dissolved in 96% ethanol. Obtained solution was placed overnight on cold (4°C) for gliotoxin crystallization [11]. The purity of gliotoxin was checked using high performance liquid chromatography as described below (3.4).

To confirm the structural correspondence of the isolated substance to glitoxin, LC-MS-ESI was carried out. Chromatographic separation and registration of mass spectra were performed on Bruker Elute UHPLC chromatograph (Bruker Daltonics, Bremen, Germany) connected to a Bruker Maxis Impact II mass spectrometer (Bruker Daltonics, Bremen, Germany) (Supplementary file).

### Minimal inhibitory concentration assay

Determination of minimal bactericidal concentration was performed using cation-adjusted Mueller-Hinton II broth (MHB; Becton-Dickinson, Sparks, MD, US). Bacteria at 10^6^ CFU/mL were incubated in 96-well microtiter plates (Eppendorf, Hamburg, Germany) containing growth media and various concentrations of gliotoxin in series of two-fold dilutions. Bacterial growth was assessed by reading and plotting the absorbance data at 620 nm obtained by the spectrophotometer Multiscan GO (Thermo Scientific, Waltham, MA, US). Wells without visible bacterial growth were plated on MHB-agar and incubated. Antimicrobial activity was expressed by the minimal bactericidal concentration (MBC), and minimal inhibitory concentrations (MIC) which was defined as the lowest antibiotic dose at which no visible growth in medium and growth on agar-plate were detected, respectively.

### Time-kill assay

Bacteria were grown to mid-log phase in Mueller–Hinton II broth (Becton-Dickinson, Sparks, MD, US). Time-kill studies were carried out based on guideline M26-A of the CLSI [12,13], using Eppendorf non-treated polystyrene 96-well plates (Eppendorf, Hamburg, Germany). The kill kinetics of gliotoxin against bacteria were tested by incubating in the medium an initial inoculum of approximately 10^6^ colony forming units (CFU) per mL with antibiotic concentrations at the 1/2 MIC, 1MIC and 2MIC. Viable cell counts were determined after 0, 1, 2, 4, 6, 8, and 24 h of incubation at 37 °C by plating serially diluted samples onto Muller-Hinton agar plates. Bactericidal activity was defined as a ≥3-log10 CFU/mL decrease, in comparison with the baseline, after 24 h of incubation.

### Absorbance of gliotoxin by bacterial cells

Evaluation of time-amount relationships of GTX absorption by the cells was performed with reversed-phase high performance liquid chromatography (RP-HPLC).

*Escherichia coli* MG 1655 and *Staphylococcus epidermidis* were cultured as described above, then pellet down by centrifugation and washed by PBS. The suspensions of each strain were diluted to 10^8^ CFU/mL and moved in separate tubes. Gliotoxin was applied in each tube and stored at 23 °C for appropriated time. The tubes that contains only gliotoxin without cells serves as negative control.

When the required time interval has elapsed, the tubes were centrifuged at 15 000 rpm for 10 min. Supernatants had been collected and were analyzed immediately (test samples).

Analytical RP-HPLC was performed using Luna C18 analytical column (4.6 × 250 mm, 5 μm, Phenomenex, US). The elution was carried out using solvent B (80% acetonitrile in ultrapure water with 0.1% TFA) in a linear gradient according to the following scheme: 0–5 % for 5 min, 5-10 % for 5 min, 10 % for 5 min, 10-20% for 20 min, 20-70% for 5 min at a flow rate of 1.0 ml/min. Absorbance was detected at 220 nm.

Chromatograms were processed using OpenLab CDS ChemStation software, and square of peak corresponding to GTX was recorded. Amount of non-absorbed GTX was calculated as the difference between the square of GTX-peaks of negative control and test sample.

### Bioluminescence assays

Heat shock response and reactive oxygen species production were assessed using various *E. coli* MG1655 bioluminescent reporter strains. Reporters carry the hybrid plasmids in which genes *luxCDABE* of the soil luminescent bacteria *Photorhabdus luminescens* are located under the control of corresponding inducible promoters *PibpA* (heat shock) or *PkatG* (hydrogen peroxide) [14].

The bacterial strain was cultivated in LB-broth medium containing ampicillin (100 μg/mL). The overnight cultures of the biosensors were washed and re-suspended in pure water to obtain a cell density corresponding to 10^7^ CFU/mL.

Ninety microliters of bacterial suspension were mixed with an appropriate volume of two-fold dilutions of gliotoxin and placed into wells of a 96-wells microplate with non-transparent side walls (Eppendorf, Hamburg, Germany) up to the final volume of 100 μL.

To assess the Trolox on the development of oxidative stress, bacterial cells were pre-incubated for half-hour with Trolox (Sigma-Aldrich, US) up to the final concentration of 400 μM.

Then, wells filled with sterile deionized water and containing appropriate amount of bacterial biosensor were used as the negative controls. Wells containing biosensor and hydrogen peroxide or ethanol were used as positive controls for oxidative stress, and folding stress, respectively

Bioluminescence measurements were carried out using the plate reader Fluoroscan Ascent FL (Thermo Scientific, Waltham, MA, US) which dynamically registered the luminescence intensity of the samples for 2 h and were estimated in relative light units (RLU). The resulting values of bioluminescence were estimated in induction factor (*R*), which was processed according to the following Equation (1):

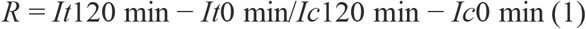

where *It* is intensity of bioluminescence of a treated sample; *Ic* is intensity of bioluminescence of the control sample (in the absence of inductor).

### Determination of lipid peroxidation

Formation of malondialdehyde (MDA) was used to measure lipid peroxidation. MDA was quantified based on its reaction with thiobarbituric acid (TBA) to form a pink MDA-TBA adduct [15].

One hundred microliters of bacterial suspensions were mixed with 200 μl of 10% (wt/vol) trichloroacetic acid, and the solids were removed by centrifugation at 10000 × *g* for 20 min. Three hundred microliters of a freshly prepared 0.67% (wt/vol) TBA (Sigma-Aldrich, USA.) solution was then added to the supernatant. The samples were incubated in a boiling water bath for 15 min and cooled, then the absorbance at 532 nm was measured with a Multiscan Go spectrophotometer (Thermo, US).

### Microscopy investigation

Bacterial cells were grown in LB-broth (Becton-Dickinson, Sparks, MD, US) to mid-Log phase, then centrifuged (15000 rpm for 10 min), re-suspended in HEPES buffer (10 mM) and adjusted to an optical density corresponding to 10^7^ CFU/mL. GTX was added to *E. coli* MG1655 and *S. epidermidis* suspensions at concentrations corresponding to their MIC.

After 4-h incubation at 25 °C the cells were washed with distilled water and stained with LIVE/DEAD BackLight bacterial viability kit (Thermo scientific, USA). Fluorescence microscopy of stained bacteria was performed using a Zeiss Axio Imager M2 fluorescent microscope (Zeiss, Oberkochen, Germany) equipped with filter sets for simultaneous viewing of Syto 9 and PI fluorescence.

### Proteins Aggregation Assay

Assay was performed according to [16]. *E. coli* MG 1655 was grown in 2.5 ml of LB medium to mid-Log phase. Bacterial cells were harvested by centrifugation at 15000 rpm for 15 min, 4 °C. The pellet was washed with 15 ml of washing buffer (50mM HEPES, pH 8.0,) and resuspended in 1.5 ml of washing buffer. Cell disruption by ultrasonication (3 × 1 min), 4 °C.

Lysates were cleared by centrifugation at 15000 rpm for 20 min, 4 °C. 40 μL aliquots were treated with 10 μL GTX (6 - 100 μg/mL) for 120 min, 30 °C. Samples were centrifuged at 15000 rpm for 20 min at 4 °C. Supernatants were mixed with 10 μL of 5×SDS sample buffer.

Pellets were washed with 100 μL of washing buffer prior to resuspension in 100 μL of 1×SDS sample buffer. 2.5 μL of each sample were separated by denaturing SDS-PAGE.

### Statistical processing

The experiments were performed using two independent series with three technical replicates each. The obtained results were statistically manipulated with Origin 2021 (OriginLab Corporation, Northampton, MA, USA) software.

The Shapiro-Wilk test was used to assess the normal distribution of values. In the presence of a normal distribution, the Student’s *t*-test has been used, indicating the mean and standard deviation (mean ± SD). Differences were considered significant at *p*-value <0.05.

## Results

The spectrum of antimicrobial activity of GTX included various types of Gram-positive and Gram-negative bacteria (Table 1). It was found that bacteria with Gram-positive cell structure died at lower concentrations, comparing to Gram-negative ones. (Table 1). MIC-values for Gram-positive and Gram-negative species were less than, and more than 10 μg/mL, respectively. Pseudomonas strains were the most resistant to GTX, while *S. epidermidis* and *C. violaceum* were the most susceptible. The MIC and MBC values in the case of all tested bacteria were the same, which characterized the mechanism of action of gliotoxin as bactericidal.

**Table 1.**
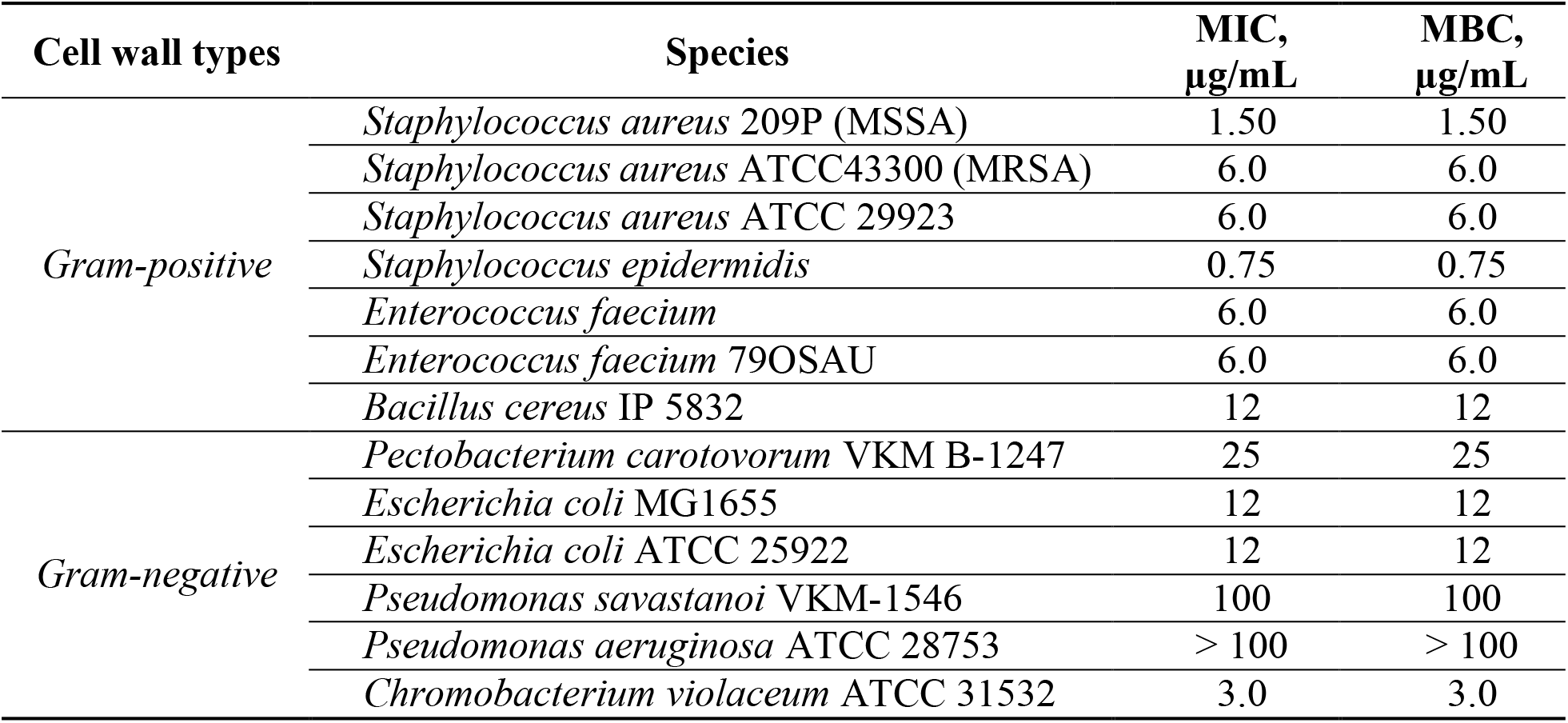
Spectrum of antimicrobial activity of gliotoxin.

### Uptake of gliotoxin

Using an analytical tool, we assessed how quickly gliotoxin will be absorbed by bacterial cells with various cell wall structures. It was found, that for the first ten minutes GTX uptake rates were 14.96±6.30 and 11.99±0.47 % for *S.epidermidis* and *E.coli*, respectively (Figure 1). During the next 50 minutes, *S.epidermidis* absorbed 37 % of gliotoxin, while the maximum absorption capacity of *E.coli* MG1655 was 45 % and have been reached at 4-th hour (Figure 1). It is need to not, that some amount of initially absorbed gliotoxin has been moved out from *S.epidermidis* at 120-th minute (Figure 1).

**Figure 1.**
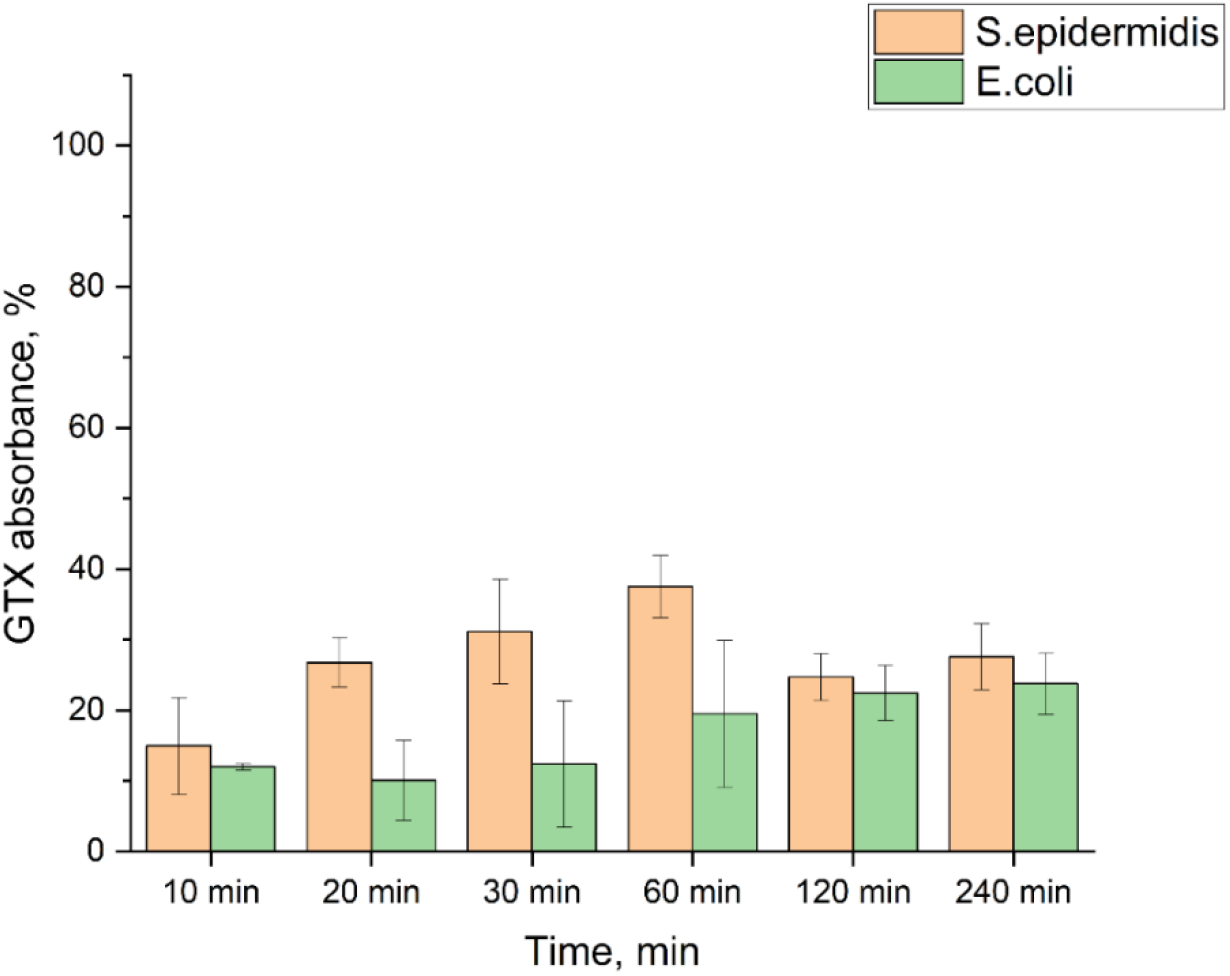
Uptake of gliotoxin (10 μg/ml) from the medium by the Gram-positive and Gram-negative bacteria. The remain amount of gliotoxin in the medium was determined by RP-HPLC and calculated by comparing the peak square of gliotoxin in medium with and without bacteria.

### Gliotoxin kills bacteria for 4 hours

The assay for the rate of bactericidal effect showed that the effect of gliotoxin does not depend on the cell wall structure, but rather depend on strain differences. Comparing the dynamics of bacterial death under the treatment, it was found that *E. coli* and *S. epidermidis* reduced its CFU value for 99 % at 4th hour of co-incubation (Figure 2 a, b, respectively). In the same conditions, bacterial cells of *S.aureus* 209P had died to 24^th^ hour (Figure S2).

**Figure 2.**
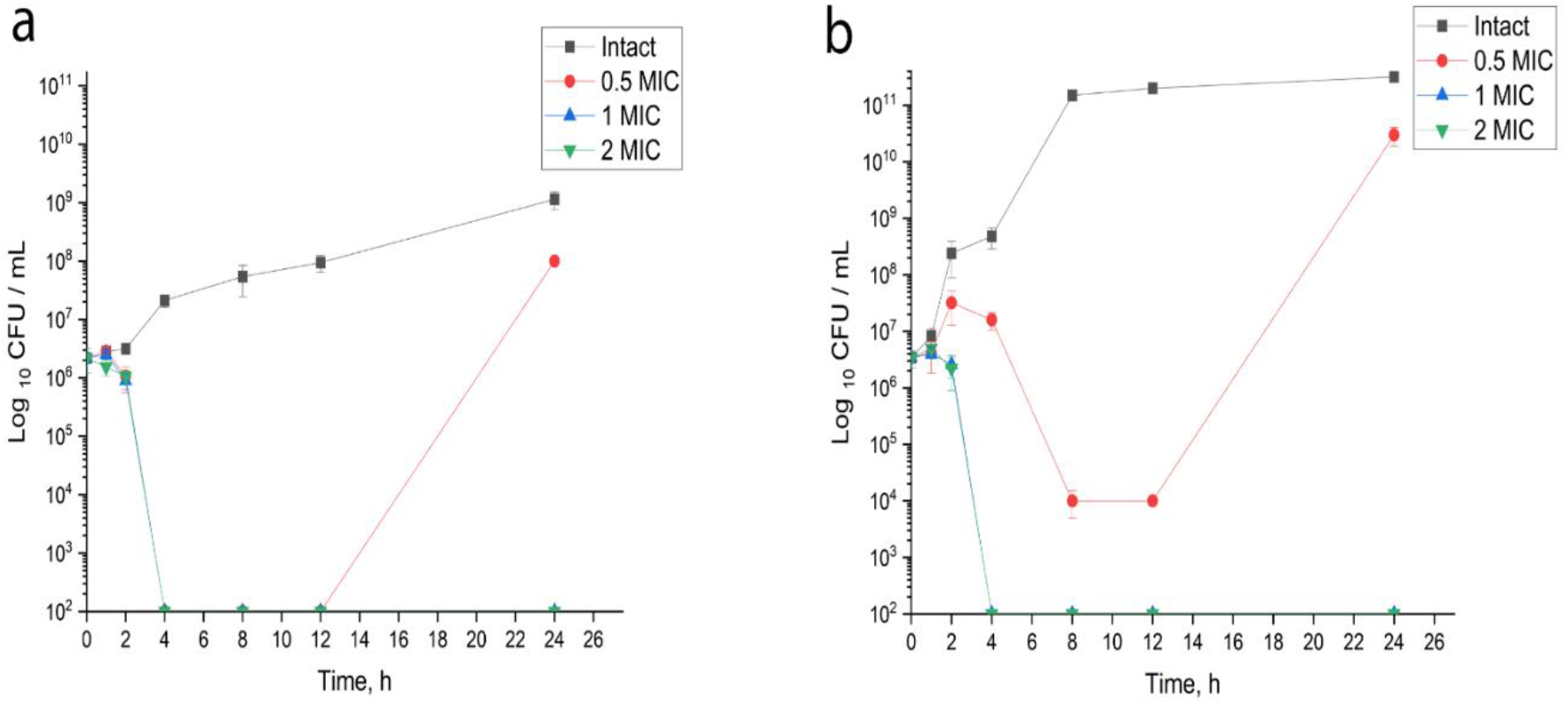
Time-kill assay with *E. coli* MG1655 (a) and *S. epidermidis* (b). The bactericidal effect was tested with gliotoxin taken at concentrations 0.5×MIC, 1×MIC, and 2×MIC in Mueller–Hinton medium.

### Gliotoxin induces oxidative stress response in bacterial cells

One of the proposed mechanisms for the antimicrobial action of gliotoxin is the generation of reactive oxygen species in the Fenton cycle. Gliotoxin’s MIC value has been re-evaluated for bioluminescence assay, since an order of magnitude more cells (10^7^ CFU/ml) was needed.

It was found that GTX in concentrations equal MIC and below (50-100 μg/ml) reduced the bioluminescence due to its toxic action. At the same time, GTX taken at sub-MIC (25-1.5 μg/ml) activated bioluminescence response with its maximum in the range 1/4-1/8 MIC (Figure 3 a). At the same time, hydrogen peroxide taken for 1/256 MIC (2.5 10^−7^ M) increased bioluminescence for 3-times higher than GTX.

**Figure 3.**
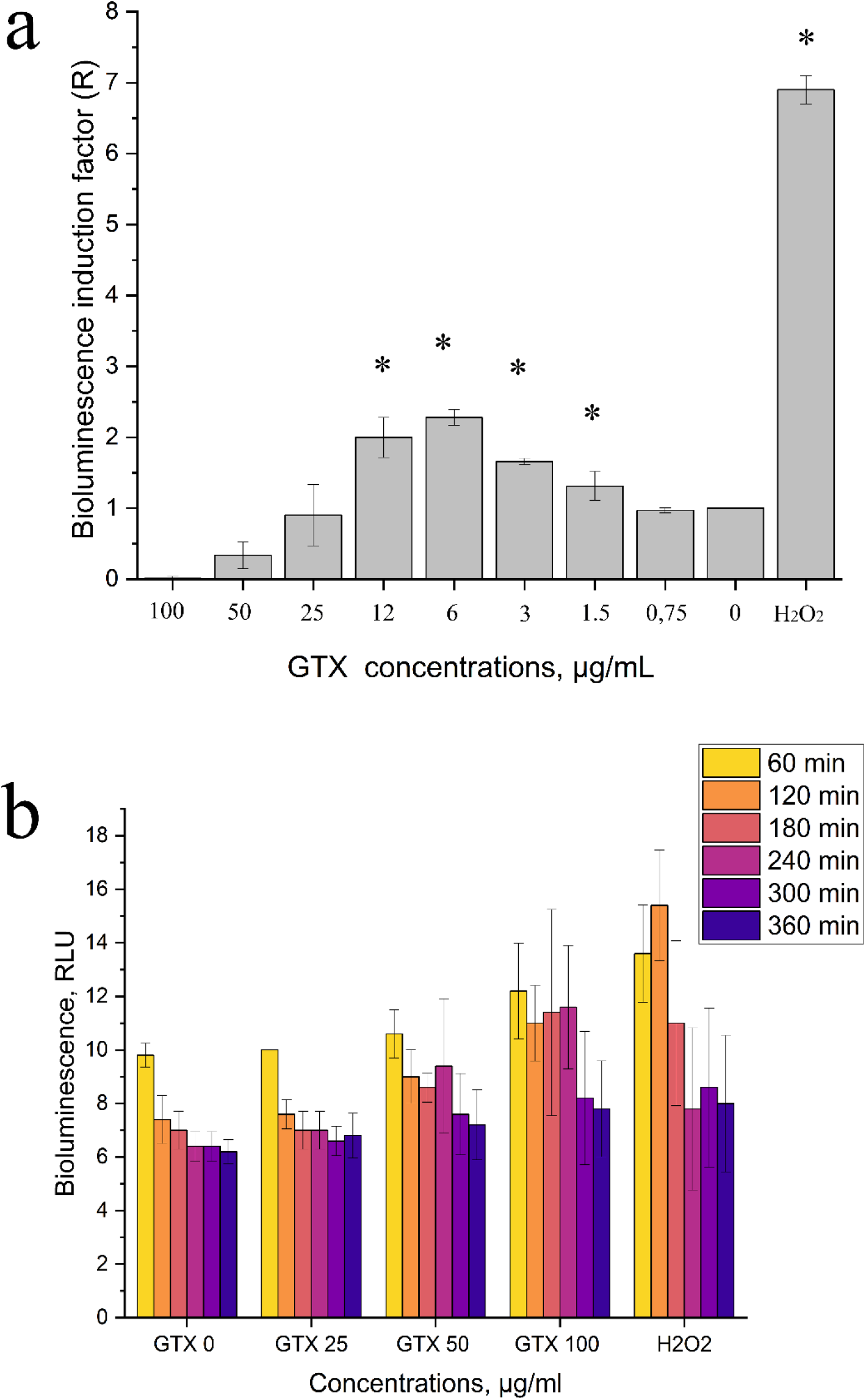
Bioluminescence response of *E. coli* MG1655 (p*KatG*’::*lux*) on gliotoxin treatment. The data presented as factor of bioluminescence induction (*R*), which is proportional to ROS concentration (a), and kinetics of bioreporter bioluminescence assayed by microchip techonologue (b). Hydrogen peroxide was taken as positive control for 2.5 10^−7^ M. Asterisks indicate a *p*-value below 0.05 as determined by an unpaired Student’s *t* test.

At the same time we designed a mirror-bottomed, less then 10 μl per well microaray made out of black polyethilene and used it for multi-well bioluminescent evaluation of GTX using gel-immobilzed cells. GTX and H_2_O_2_ induced bioluminescent response is concentration-depended as aspected, maximized at 60-120min incubation time and slowly faded aftewards (Figure 3 b). The microwell arrays approach provide an orded of magnitude more data per surface unit and use an orded of magnitude less reagents, so it will substitute classical plates in our future work.

Involving of reactive oxygen species in the bactericidal action of gliotoxin was evaluated using antioxidant Trolox. It was found that Trolox (400 μM) completely inactivated the bioluminescence of the biosensor that is evidence that oxidative stress is absent. At the same time, addition of Trolox did not change MIC values of gliotoxin.

Thus, it was found that gliotoxin does indeed generate reactive oxygen species in bacterial cells, but only slightly above the natural level. Thus, ROS is unlikely the main mechanism of GTX antibacterial action.

### Gliotoxin does not oxidize cells lipids

The formation of ROS in cells leads to the oxidation of various molecules, in particular, phospholipids of the cytoplasmic membrane and cell wall. Determination of peroxidation of malonic aldehyde was carried out in the presence of thiobarbituric acid. Solution of hydrogen peroxide was taken as a positive control caused lipid peroxidation in the MIC-range (Figure 4 a). It was found that GTX at bactericidal concentrations slightly peroxidase cell’s lipids. However, the differences in peroxidation were insignificant from control values (intact cells) (Figure 4 b).

**Figure 4.**
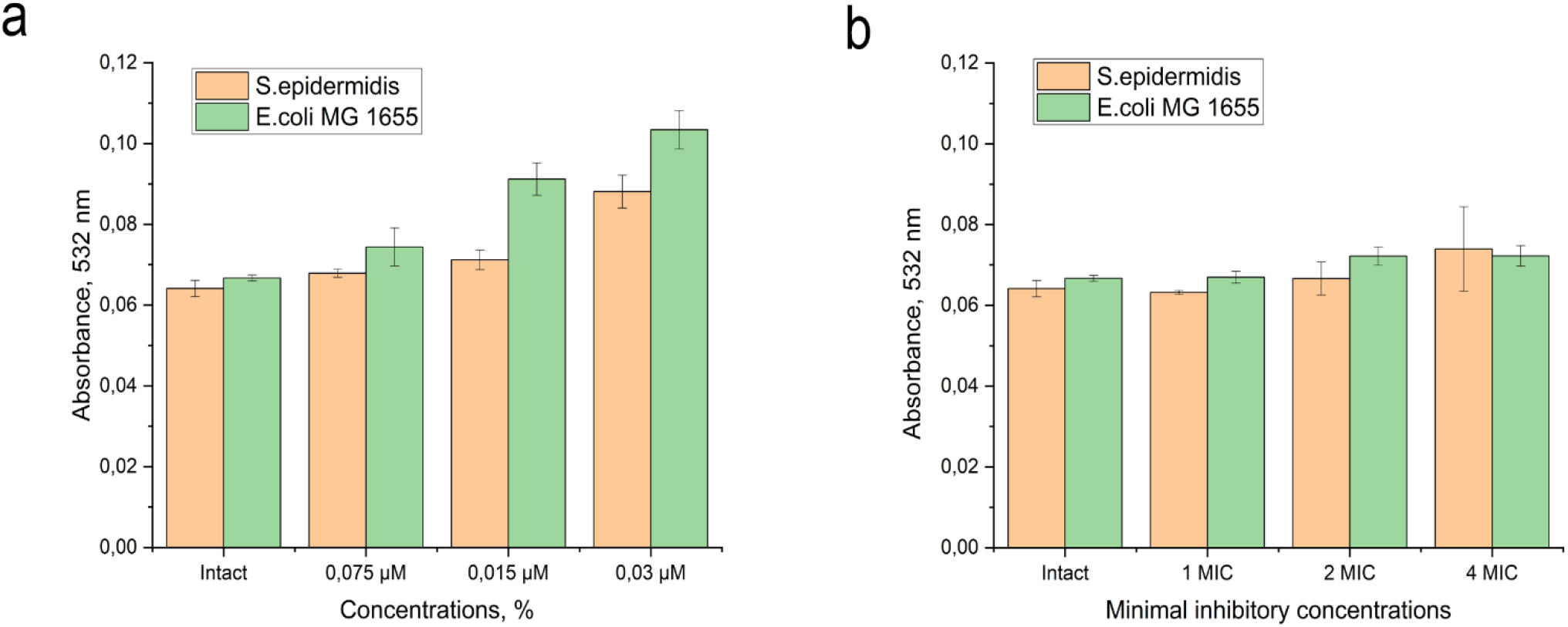
Effect of gliotoxin on lipid peroxidation of *E.coli* MG 1655 p*KatG’::lux* and *S.epidermidis*. a) Hydrogen peroxide was taken as positive control: 1/8 MIC - 1 MIC (2×10^−5^ – 3×10^−3^ M). b) Bacterial cells were treated with gliotoxin at concentrations corresponding to 1-4 MIC (50-200 μg/mL) for 4 hours.

### Gliotoxin disturbs bacterial cell wall

Fluorescence microscopy was used to evaluate the effect of GTX on bacterial cell membranes. It turned out that treatment with gliotoxin for 4 hours led to loosing of structural integrity of bacteria.

The obtained images show that in the control samples of *E. coli* and *S.epidermidis* contain significant part of the cells in the population with intact membrane (Figure 5 a, c, respectively). Treatment of cells with gliotoxin at a concentration of 1MIC lead to the accumulation of cells with damaged barrier structures (fluoresce in the red range) (Figure 5 b, d, respectively).

**Figure 5.**
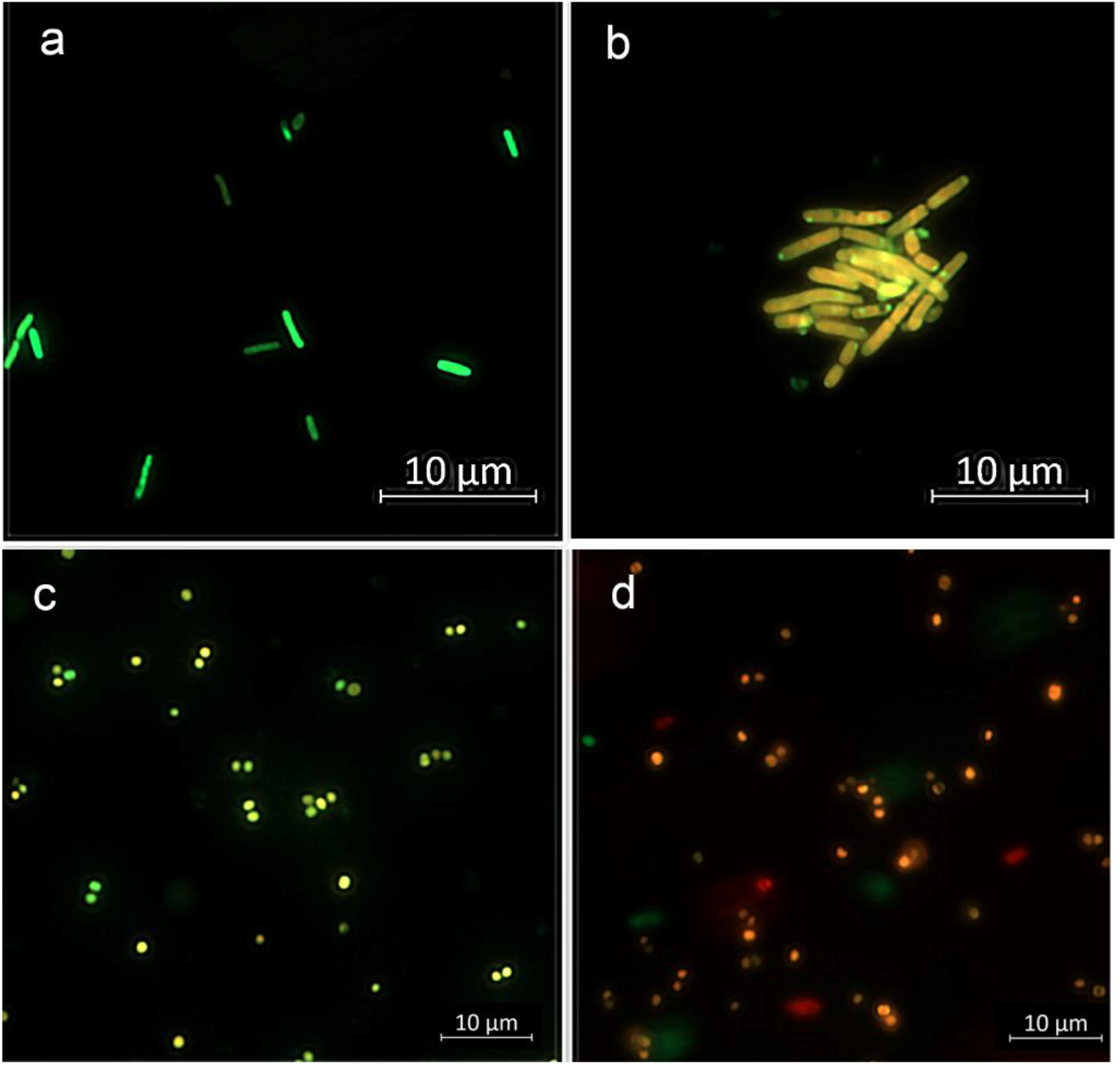
Assessment of the structural integrity of cell membranes under the influence of gliotoxin. Epifluorescence images of untreated *E.coli* MG 1655 (a) and *S.epidermidis* (d) cells, and treated with GTX (b, d).

Thus, the action of gliotoxin affects the integrity of cellular barrier structures.

### Thiol-containing substances inhibit antibacterial action of gliotoxin

It is known that molecules with disulfide bonds, such as allicin and gliotoxin, can interact in animal cells with free thiol groups of intracellular proteins. If this is true, then the introduction of various thiol-containing substances into the reaction medium should reduce the antimicrobial activity of gliotoxin. We modified the MIC-assay by adding to the medium 2-mercaptoethanol (2-ME), reduced glutathione (GSH), and Trolox. It turned out that addition of antioxidant Trolox does not affect the bactericidal properties of gliotoxin, while 2-ME and GSH increase the MIC values by eight times (Table 2).

**Table 2.** Effect of Thiol-Containing substances and antioxidant on the antimicrobial activity of gliotoxin, expressed in terms of the minimum inhibitory concentration

| MIC, μg/ml | Gliotoxin, μg/mL | Trolox 400 μM | 2-mercaptoethanol | Reduced glutathione, 5 mM |
| --- | --- | --- | --- | --- |
| *S.epidermidis* | 1.0 | 1.0 | 25.0 | 25.0 |

Thus, modification of free cysteine thiols is a cause for bacteria inhibition, since addition of various thiols promote bacterial surviving.

### Gliotoxin causes protein aggregation and heat shock response

It is known, that sulfur-containing antibiotics like as allicin induce modification of proteins unfolding and aggregation leading to unfolding stress [16]. To evaluate the ability of gliotoxin to cause unfolding stress, we evaluated aggregation of proteins by GTX treatment. After the treatment, cells proteins were divided by centrifugation on soluble and insoluble fractions, and separated by SDS-PAGE electrophoresis (Figure 6 a).

**Figure 6.**
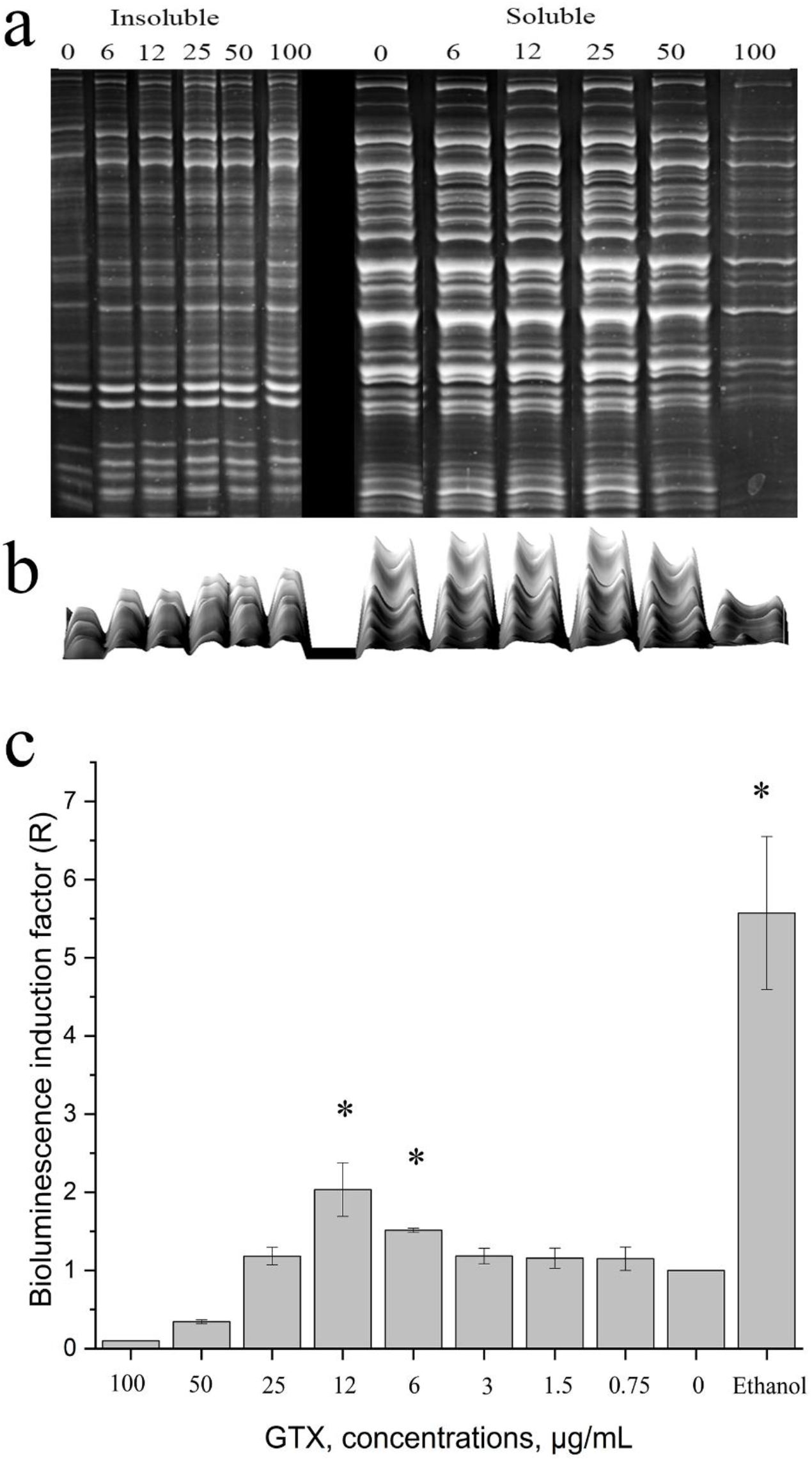
Gliotoxin induced protein aggregation and unfolding stress. Crude extract from *E.coli* MG1655 was treated with different concentration of GTX for 15 min at 25 °C. Proteins were separated into soluble and insoluble fractions by centrifugation prior to SDS-PAGE. Results were processed by ImageJ software, there intensicity of protein bands were imaged in pseudo-3D manner (b).

It was found protein aggregation in a concentration-dependent manner, that was already visible at 25 μg/ml (Figure 6 b). Using a biosensor strain we assayed whether the proteins aggregation by gliotoxin caused heat shock response. Biosensor *E.coli* MG 1655 carries the hybrid plasmids in which cassette of *luxCDABE* genes are under the inducible promoter *PibpA* (heat shock). It was found slightly but statistically significant activation of bioluminescence by addition of gliotoxin at sub-MIC concentrations (Figure 6 c).

Bioluminescence response of *E. coli* MG1655 (p*IbpA*’::*lux*) on gliotoxin treatment (c). The data presented as factor of bioluminescence induction (*R*), which is proportional to unfolding stress. Water solution of ethanol (4.5 %) was taken as positive control. Asterisks indicate a *p* value below 0.05 as determined by paired Student’s *t* test.

Our data indicate that gliotoxin cause protein aggregation *in vitro* and induce the heat shock response *in vivo*.

## Discussion

The first study of the antibacterial action of gliotoxin refers to the first half of the 20th century [11, 17, 18]. To date, a lot of data has been accumulated on the effect of gliotoxin on various eukaryotic cells [15]. Data on gliotoxin antimicrobial properties are accumulating [19–21], which allows gliotoxin to be considered in antibiotic monotherapy or in combination [18]

Existing studies had shown very quickly uptake rate of gliotoxin by various mammalian cells, most of the toxin becomes cell associated within 10 min [22]. In our work we have found significant decreasing in free gliotoxin during first 10 minutes, which is consistent with the data obtained on fungi [23].

Absorption of gliotoxin did not lead to immediate cell death. It takes at least 4 hours for decreasing the number of live bacterial cells by 99.99%. It is interesting, that gliotoxin action against Gram-negative and Gram-positive bacteria was differ in concentrations, but the same in killing time. In the case of Gram-negative bacteria, few times more gliotoxin was required. The main explanation for this is that Gram-negative bacteria have an outer membrane that contains significantly more proteins than the cell wall of Gram-positive bacteria. [24]. Earlier, Jones and Hancock showed that *Salmonella typhimurium*, deficient in outer-membrane polysaccharide synthesis, was hypersensitive to gliotoxin. Thus, gliotoxin attacks Gram-positive bacteria more effectively than Gram-negative.

The presence of a disulfide bridge in the epipolythiodioxopiperazine (EPT) molecules makes it possible for several events to occur in cells at once. As a result of the generation of reactive oxygen species during the conversion of the oxidized form of EPT into the reduced form, DNA is damaged, lipids and proteins are oxidized. According to this mechanism, the toxic effect of sporidesmin is realized [1]. ROS induction as the main mechanism of gliotoxin toxicity has been established in several studies and it has been found that GTX caused single-stranded and double-stranded break of plasmid DNA *in vitro* [8]. Another work described that DNA repair-proficient strain *Escherichia coli* WP2 *trp*E56 was more resistant to GTX action, than its repair deficient derivative [25]. However, the Ames *Salmonella* assay and SOS-chromotest do not support the repair assay results. And there are several others evidence concerning to DNA-reactivity of gliotoxin in vitro studies [8, 26, 27].

In our study we used a bacterial bioluminescent sensor for the detection of reactive oxygen species. It was found that gliotoxin induces the formation of hydrogen peroxide in bacterial cells. However, concentration of formed H_2_O_2_ was insufficient for lipid peroxidation, as we found. To clarify involvement of oxidative stress in toxic effects of gliotoxin, we pre-incubated bacterial cells with Trolox, a vitamin E analogue. Earlier, free radical scavengers including vitamin E have been shown to inhibit the toxic effects of another EPT sporidesmin [28]. We have shown that pre-incubation of bacterial cells with Trolox reduced bioluminescent response, but did not reduced bactericidal properties of gliotoxin. These results are in agreement with the study where radical inhibitors failed to prevent apoptosis in thymocytes induced by gliotoxin [29]. Collectively, these results indicate that formation of reactive oxygen species is not significant for the bactericidal activity of gliotoxin.

The more likely that GTX interacts with cells proteins. Gliotoxin has been shown to inhibit alcohol dehydrogenase by interaction with a specific thiol group (either Cys-281 or −282), with the formation of a mixed disulphide [30]. The work of Cavallito group showed that gliotoxin activity lost when treated with an excess of cysteine and thyoglycolate [31]. In our work we had found that addition of various thiol-containing substance significantly inhibited the gliotoxin’s bactericidal action. Moreover, gliotoxin treatment of crude bacterial lysate led to aggregation of bacterial proteins, but in less pronounced manner as it has been revealed with allicin [16]. The consequence of cellular protein aggregation is a heat shock response, as we found with bioluminescent reporter strain. These result is similar with allicin mode of action, that is a combination of decrease in glutathione level, unfolding stress, and inactivation of crucial metabolic enzymes through *S*-allylmercapto modification of cysteines [16].

The consequence of gliotoxin interaction with bacterial cells is disturbance of their barrier structures. Jones and Hancok assume in their work that it is possible that reaction of gliotoxin with membrane thiol groups can lead to increased membrane permeability, but they did not determine whether leakage occurred [23]. However, there is evidence that *C. albicans* yeast cells treated with gliotoxin became more permeable [32].

Thus, gliotoxin demonstrated promising activity against bacteria, and the results obtained indicate that mechanism of its antimicrobial action against prokaryotes is differ than toxic action on eukaryotes.

