## Supplementary Figures for "Antibacterial Action of Gliotoxin Realized Through Interaction with Tiol-containing Proteins"

**Gliotoxin identification by mass-spectrometry**

**Material and methods**

Chromatographic separation and registration of mass spectra were performed using Bruker Elute UHPLC chromatograph (Bruker Daltonics, Bremen, Germany) connected to a Bruker Maxis Impact II mass spectrometer (Bruker Daltonics, Bremen, Germany) with a working resolution of 20000, equipped with an electrospray ionization source. Chromatographic separation was carried out in the gradient elution mode. The mobile phase was 0.1% aqueous formic acid (eluent A) and 0.1% formic acid in acetonitrile (eluent B) at a flow rate of 0.3 ml/min. InfinityLab Poroshell reverse phase column was used for separation (2.1×150 mm, phase particle diameter 1.9 µm (Agilent Technologies, Santa Clara, CA, USA). The gradient program was as follows: 0 min - 5% B, 20 min - 100% B, 25 min - 100% B, 25.2 - 5% B and 28 min - 5% B.

Mass spectrometric detection was carried out in the positive ion detection mode in the range of 50–2000 Da. The operating parameters of the ionization source were as follows: the capillary voltage was 4500 V; nitrogen was used as a drying gas at a flow of 6 l/min, a pressure of 2.5 bar, and a temperature of 200°C. The voltage at the first and second ion funnels was 300 V, the voltage at the lenses of the quadrupole and the collision cell was 5 and 10 V, respectively. MS/MS spectra in the mode of dissociation induced by collisions were obtained automatically, nitrogen was used as a gas for collisional activation. To obtain MS/MS spectra, peaks were selected in the range of 130–1500 m/z with an intensity greater than 400 relative units. The activation energy was chosen depending on the mass of the precursor ion and varied in the range from 15 eV for 100 m/z to 100 eV for 1500 m/z.

**Results**

When separating the active fraction of metabolites *Aspergillus fumigatus* UTMN1, an ion chromatogram was obtained (Figure 1*).* The main peak was detected on the chromatogram was the peak with retention time (rt) 8.1 min. The panoramic spectrum corresponding to this peak is shown in Figure 2.

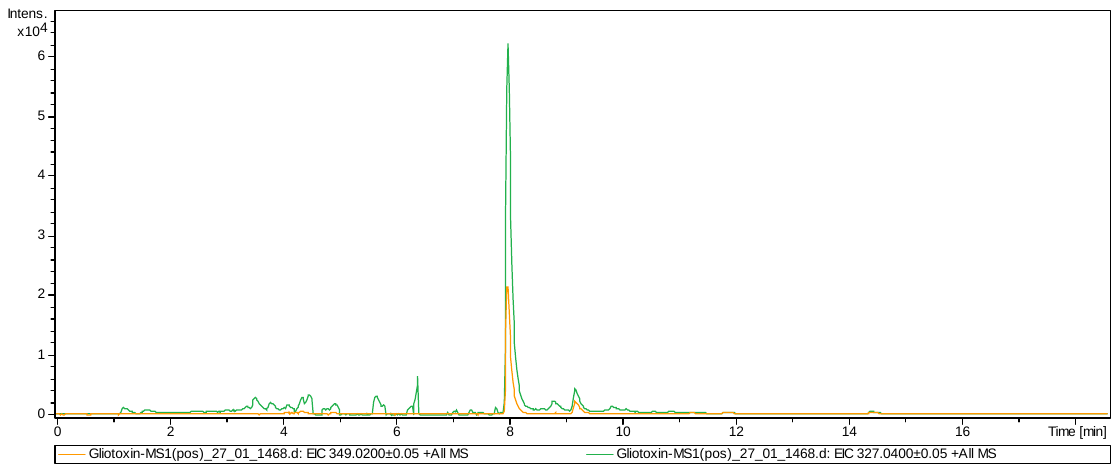

Figure S1. Chromatograms for ion current at m/z 349.02±0.05 and 327.04±0.05 corresponding to gliotoxin ions [M+H]^+^ and [M+Na]^+^ (top - sample, bottom - blank ).

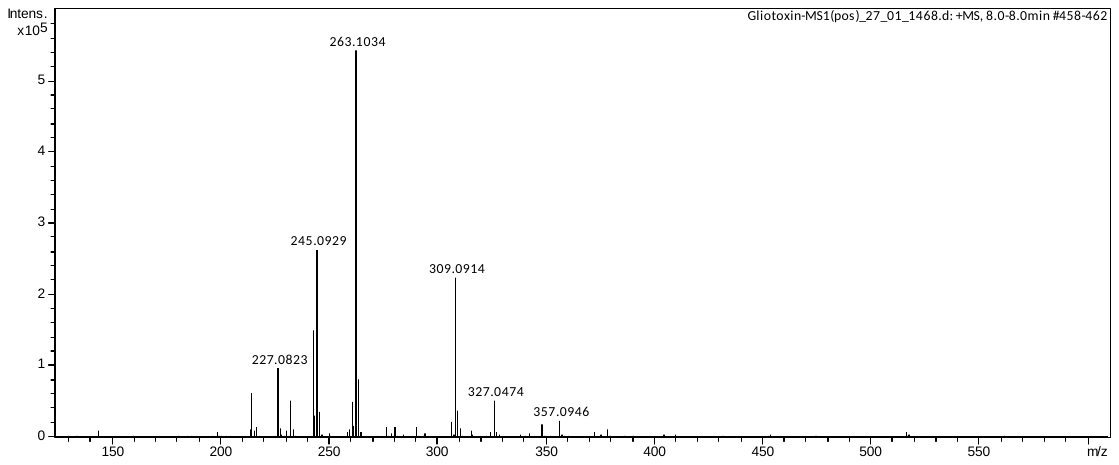

Figure S2. The panoramic mass spectrum of the main component (rt 8.1 min).

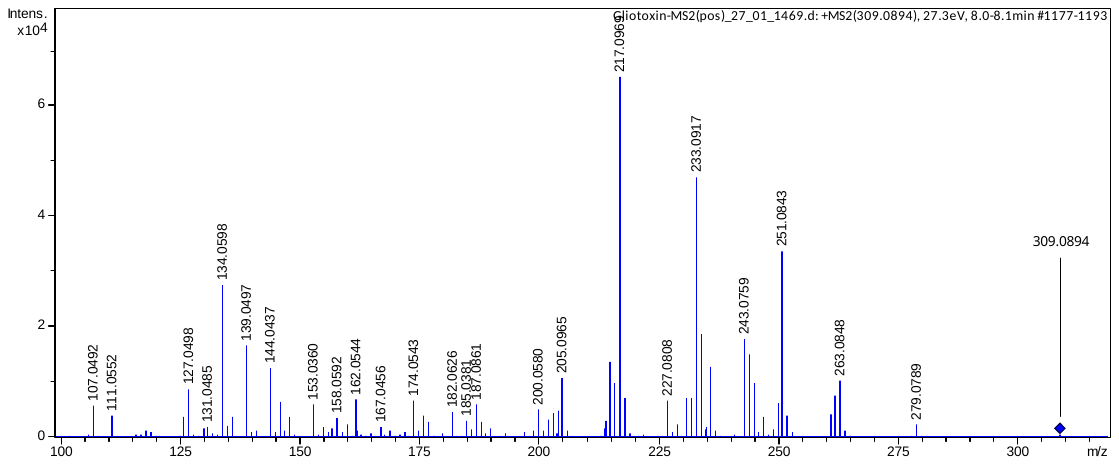

Figure S3. MS2 peak spectrum at m/z 309.09 (rt 8.1 min).

The brutto-formula C_13_H_15_N_2_O_4_S_2_ and value m/z 327.0474 which were calculated are correspond to gliotoxin (PubChem ID 6223) (Table 1).

Table S1. Calculated parameters of the main component (rt 8.0 min).

| Meas. m/z | Ion Formula | m/z | err [ppm] | mSigma | rdb | e? Conf | N-Rule |
| --- | --- | --- | --- | --- | --- | --- | --- |
| 327.0474 | C13H15N2O4S2 | 327.0468 | -2 | 17.9 | 8 | even | ok |

**Estimation of gliotoxin mode of action using microchip technique**

**Material and methods**

The microplate was made by milling an aluminum composite panel (fig.1). The agarose solution was brought to a boil and cooled at room temperature to 40°C. Then mixed with the culture of *E. coli* MG 1655 pKatG-lux in a ratio of 1:1. The resulting mixture was incubated for 5 minutes and added to the wells of the tablet for 2 µl. 0.2 µl of gliotoxin was added to the resulting mixture at concentrations of 0-100 µg/ml. As a positive control, 0.03% hydrogen peroxide was added instead of gliotoxin. The kinetics were filmed with a RisingTech CCD camera (Model: ATR3). Exposure of 1 frame was 15 minutes. Data were quantified using a self-written C Sharp script. The intensity of luminescence (relative light units, RLU) was estimated as the sum of the brightness of all pixels in the selected area divided by the total number of pixels.

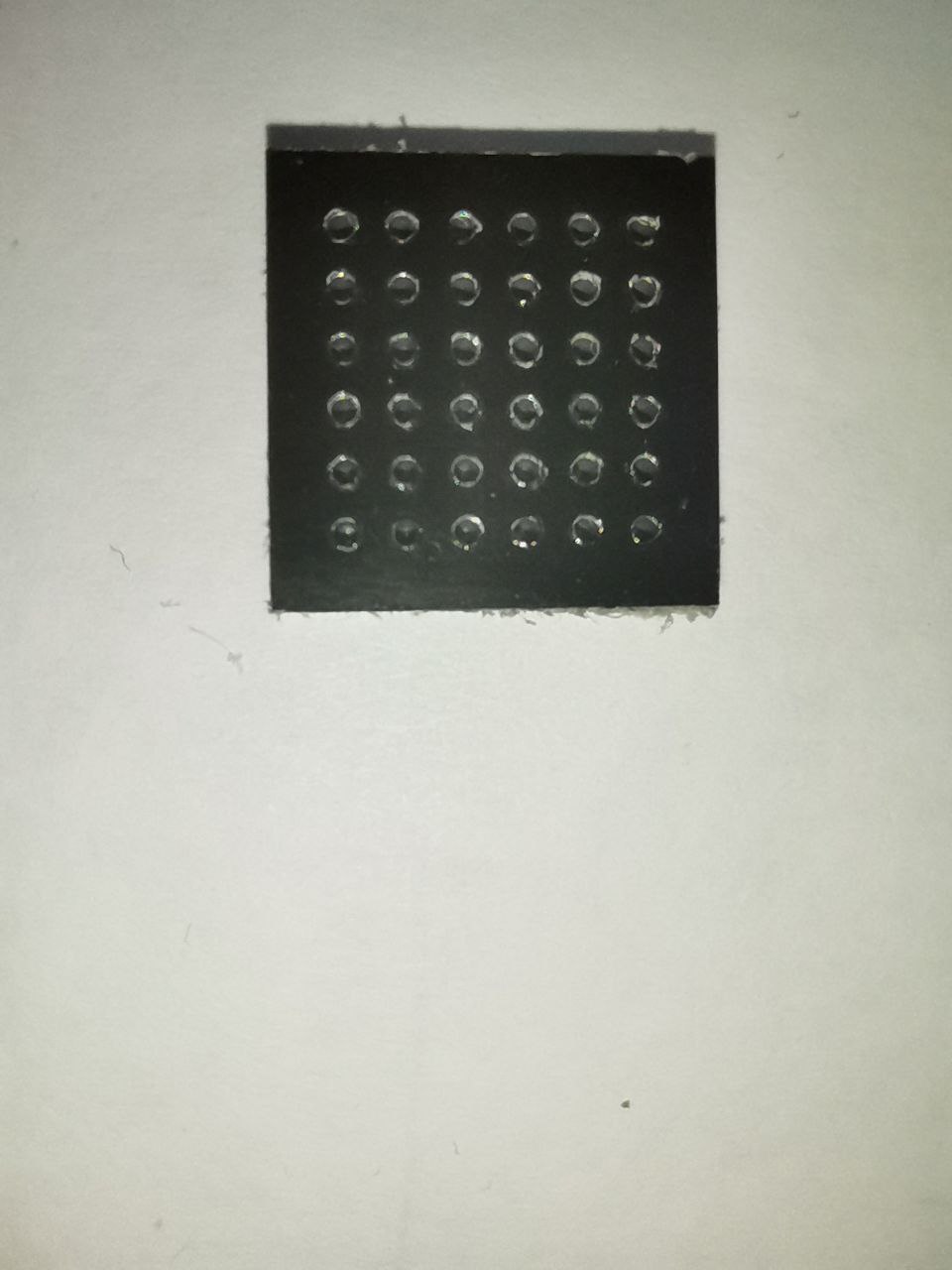

Figure S4. Aluminum composite panel: enamel, aluminum 300µm, PE 3mm, aluminum 300µm, enamel

**Results**
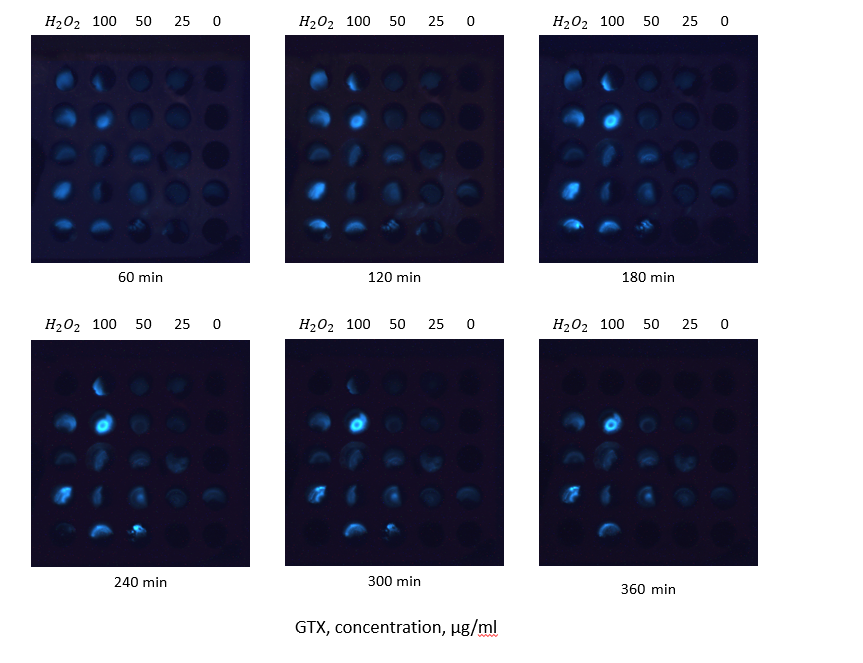

Figure S5. Kinetics of *E.coli* MG 1655 pKatG-lux bioluminescence under the treatment with gliotoxin. The frames have been captured in microchip wells (volume – 2.2 µl) using CCD camera at various time-intervals., hydrogen peroxide - 0.03%, Gliotoxin concentrations – 0-100 µg/ml.
